# Deciphering Tumour Tissue Organization by 3D Electron Microscopy and machine learning

**DOI:** 10.1101/2021.06.15.446847

**Authors:** Baudouin Denis de Senneville, Fatma Zohra Khoubai, Marc Bevilacqua, Alexandre Labedade, Kathleen Flosseau, Christophe Chardot, Sophie Branchereau, Jean Ripoche, Stefano Cairo, Etienne Gontier, Christophe F. Grosset

## Abstract

Despite recent progress in the characterization of tumour components, the tri-dimensional (3D) organization of this pathological tissue and the parameters determining its internal architecture remain elusive. Here, we analysed the spatial organization of patient-derived xenograft tissues generated from hepatoblastoma, the most frequent childhood liver tumour, by serial block-face scanning electron microscopy using an integrated workflow combining 3D imaging, manual and machine learning-based semi-automatic segmentations, mathematics and infographics. By digitally reconstituting an entire hepatoblastoma sample with a blood capillary, a bile canaliculus-like structure, hundreds of tumour cells and their main organelles (e.g. cytoplasm, nucleus, mitochondria), we report unique 3D ultrastructural data about the organization of tumoral tissue. We found that the size of hepatoblastoma cells correlates with the size of their nucleus, cytoplasm and mitochondrial mass. We also discovered that the blood capillary controls the planar alignment and size of tumour cells in their 3D milieu. Finally, a set of tumour cells polarized in the direction of a hot spot corresponding to a bile canaliculus-like structure. In conclusion, this pilot study allowed the identification of bioarchitectural parameters that shape the internal and spatial organization of tumours, thus paving the way for new investigations in an emerging field that we call ‘onconanotomy’.

## INTRODUCTION

Diagnostic, prognostic and predictive clinical cancer studies are mainly based on the analysis of tissue sections in two dimensions (2D). However, tumours often present a complex architecture that 2D imaging cannot capture. Tumour cells establish composite interactions with the surrounding cancerous cells, extracellular matrix and stromal components, including blood capillaries and immune cells. In recent investigations, we noted that various hepatoma cells engrafted on the chick embryo chorioallantoic membrane (CAM) were arranged histologically and microscopically in a lineage-specific way while growing in a comparable controlled environment ^1,2^. Moreover, pathological tumour atlases display a multitude of 2D images with tumour tissues whose cells are organized differently while belonging to the same category of cancer (http://www.pathologyatlas.ro; https://www.proteinatlas.org; https://atlases.muni.cz). These observations suggest that the structural organization of tumoral tissue changes from one sample to another and is likely controlled by environmental and genetic factors such as the tissue origin, state of differentiation, genetic programme, the mutations they harbour, the activated molecular pathways, and the surrounding stromal components such as blood capillaries. Moreover, these interactions are dynamic and vary as the cancer progresses and metastasizes. To shed light on this critical issue, we decided to investigate the ultrastructural pattern of cancer tissue.

The applications of 3D electron microscopy (EM) are still under development. Since its advent in the 1930s, EM has allowed the in-depth analysis of a wide range of biological samples. Transmission electron microscopy (TEM) and scanning electron microscopy (SEM) are routinely used to collect ultrastructural data from biological specimens at nanoscale resolution and to correlate structural images with biological functions. Although effective, these techniques produce images only in two dimensions. Therefore, investigating a large volume (100-1000 pL) of biological tissue in 3D at a high resolution was hardly possible until the development of 3D EM technologies including serial block-face (SBF)-SEM ^3^ and Focus Ion Beam-SEM ^4^.

Hepatoblastoma (HB) is the most common form of liver cancer in young children. Recently, we reported the classification of these paediatric tumours in three groups, the use of Velcade^®^ as a new therapeutic option for the treatment of aggressive HB and the development of an HB model in chick embryo for biological studies ^1,5,6^ Importantly, we developed HB-patient-derived xenografts (PDX), a tumour model that fully recapitulates the histological, genetic and biological characteristics of parental HB ^7^. Here, we used SBF-SEM to investigate the 3D internal organization of HB samples obtained from HB PDXs. Following image acquisition by SBF-SEM, we gathered quantitative data about the ultrastructure of tumoral tissue using 3D imaging tools, mathematical algorithms and deep-learning approaches. We accurately measured the size of tumour cells and their main subcellular components (e.g. nucleus, mitochondria, cytoplasm), their planar alignment, their spatial orientation and the distance between the cells and a blood capillary. Results validate the relevance of our integrated workflow using wide-field 3D EM and computational approaches to investigate the deep internal architecture of HB tissue and the spatial organization of the tumour cells. These structural and functional data pave the way for future investigations in the emerging field of onconanotomy.

## RESULTS

### Image acquisition by EM and SBF-SEM of HB PDX tissues

To study the 3D organization of HB-PDX tissues, we adapted the protocol described by Deerinck et al. ^8^ to tumour samples (Extended data Fig. 1) and analysed the ultrastructural morphology and the region of interest (ROI) of each stained sample by TEM using ultrathin sections. Fig. 1A shows that the staining of the four samples was uniform and well contrasted and that all HB tissues retained their structure of origin. For example, membranes, mitochondria, endoplasmic reticulum, nuclei, lipid droplets and circulating red blood cells into capillaries (“bc”) were clearly identifiable and typical with an excellent grey tone scale. The morphology of endothelial and immune cells was typical. Intercellular spaces were also visible and identified as bile canaliculus-like structures (“bi”, Sample 2)^9,10^, which are indicative of liver-derived tissue. Overall, these results demonstrated the preservation of the ultrastructure of our HB samples following fixation and staining procedures.

**Fig. 1:**
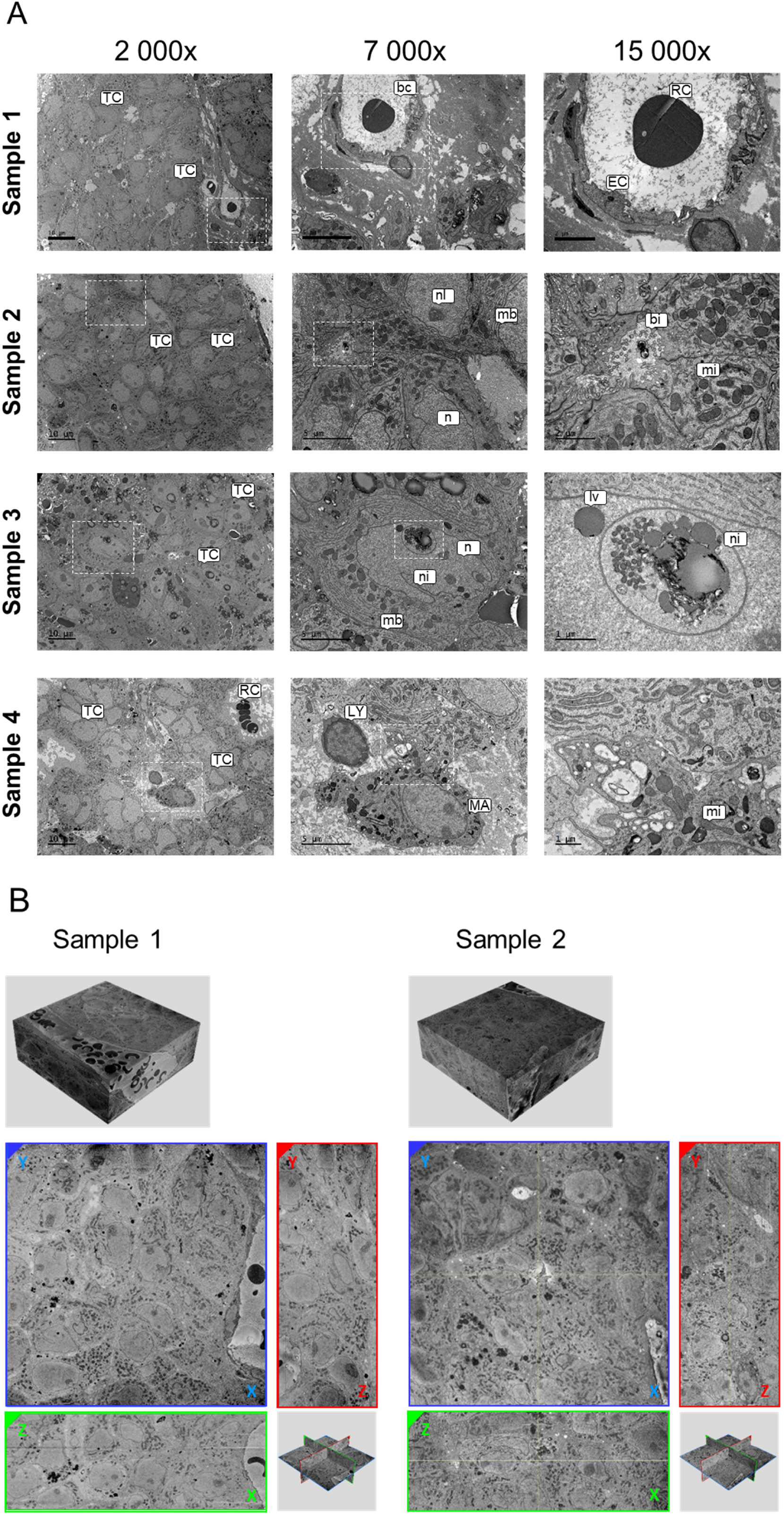
Analysis of HB-PDX by EM. (A) Analysis of four HB-PDX by TEM at different magnifications. White dotted-line frame: ROI corresponding to immediately higher magnification. bc, blood capillary; bi, bile canaliculus-like structure; EC, endothelial cells; MA, macrophage; lv, lipid vesicles; mb; plasma membrane; mi, mitochondrion; n, nucleus; ni, nuclear invagination; nl, nucleolus; LY, lymphocyte; RC, red cells; TC, tumour cell. (B) Top panels: SBF-SEM stacks from two samples. Bottom panels: Extracted images of orthogonal plane from different faces of reconstructed volume (XY, YZ, XZ), whose positions are displayed on orthoslice view positions inside stack.

Next, we analysed two HB-PDXs by SBF-SEM and acquired high-resolution images in X and Y along the Z-axis to produce a substantial volumetric view of each sample. To obtain both image quality and detailed volumetric information, the acquired volume was about 70 μm × 70 μm × 25 μm (pixel time = 10 μs, pixel size = 15 nm). After acquisition, the images were aligned to obtain 3D stacks (Fig. 1B; Sample 1: 250 images, 113.9pL; Sample 2: 246 images, 121.6pL). While volumetric image resolution was not isotropic (15 nm lateral resolution *versus* 100 nm depth resolution), ultrastructural features within the 3D stack were perfectly maintained and identifiable, allowing us to visualize the cells and their subcellular components within the tumour samples from any angle of view on the three axes (Fig. 1B).

### 3D organization of individual tissue components

Following stack reconstruction, we analysed the tissue features in tumour sample 1 (Extended data Fig. 2A-B, Supplementary Video 1). Using Ilastik^11^ and VAST-Lite^12^ software, we manually segmented a blood capillary and its circulating red and white blood cells (one leukocyte and one structure evoking a dendritic cell or a platelet cluster) (Extended data Fig. 2C-F, Supplementary Video 1). The volume of the blood capillary portion irrigating the tissue was 11.3 pL and the 11 entirely segmented murine red blood cells were 61.92 ± 16.26 μm^3^ in size (Extended data Fig. 2C-F, Supplementary Video 1). Next, we manually segmented a tumour cell from sample 2 by delimitating the cytoplasm, the nucleus and the mitochondria. Again, the 3D digital representation allowed volumetric quantification of the cell and its cytoplasm, nucleus and mitochondrial network (970.6, 657.1, 313.6 and 64.6 μm^3^, respectively), as well as affording a morphologic view of the tumour cell in its 3D environment (Extended data Fig. 3; Supplementary Video 2). The size of this tumour cell was consistent with the size range of HB cells^13,14^.

### Reconstruction of tumour 3D organization by semi-automated segmentation

Although it produces accurate data, manual segmentation is too laborious and time-consuming to allow the analysis of hundreds of cells constituting a stack of about 110-120 pL. To overcome this problem and anticipate the analysis of larger volumes of tumour (> 200 pL), we implemented a semi-automatic segmentation algorithm comprising the manual segmentation of the cytoplasm and nucleus of cells on 1 out of 10 images in the stack form sample 2 and an automatic segmentation algorithm of ROI by proximal propagation along the Z axis (Extended data Fig. 4). This procedure allowed us to segment partially and entirely 182 tumour cells, 113 nuclei, 1 immune cell and 3 portions of the same blood capillary (Fig. 2A-C, Extended data Fig. 5, Supplementary Video 3). The elongated shape of the immune cell, the presence of many vacuoles in its cytoplasm and its location near the blood capillary was suggestive of a tumour-infiltrating macrophage (Extended data Fig. 5, Supplementary Video 3). Following the alignment in a plane of 47 tumour cells with a complete nucleus, we found that 36 tumour cells (76.6%) and 35 nuclei (74.5%) had an inclination angle of 0 to 20° (relative to the best alignment plane, see Online Methods), while the inclination angle of the blood capillary was 8° (Fig. 2D-E, Extended data Fig. 6, Supplementary Video 4). These data suggested that the blood capillary may influence the spatial arrangement of tumour cells.

**Fig. 2.**
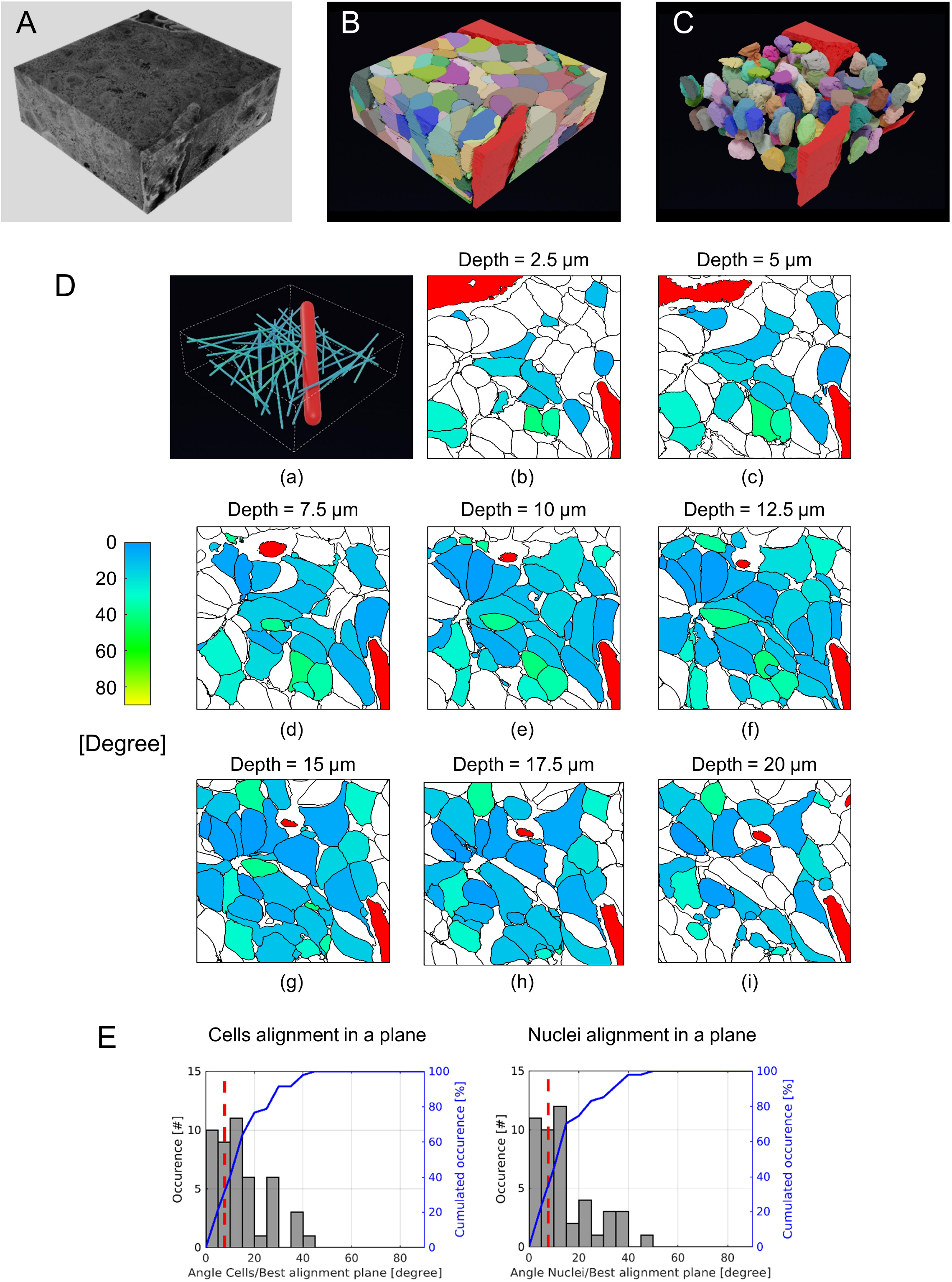
Alignment of HB cells and nuclei. (A) Sample 2 stack. (B-C) Digital representation of A with 182 cells (B) and 113 nuclei (C). (D) Alignment of cells and blood capillary in a plane: (a) coloured sticks = main axis of tumour cells; red stick = main axis of blood capillary. (b-i) Angles between each main cell axis and the best alignment plane. 2D cross-sectional maps for increasing depth along Z-axis. (B-D) Blood capillary portions are in red. (E) Histograms of cells (left panel) and nuclei (right panel) alignment angles. Red dashed line: alignment angle of blood capillary.

Next, we measured the tumour cell polarity using a vectoral approach (see Online Methods). In both cases, data showed that a set of tumour cells polarized in the direction of a hot spot corresponding to a bile canaliculus-like structure (Fig. 3A-B, Extended data Fig. 7, Supplementary Video 5)^9,10^. Numerous membrane protrusions were visible at the boundary between the bile canaliculus-like structure and the canalicular membrane of the surrounding tumour cells (Extended data Fig. 7, Supplementary Video 5). This structure into which hepatocytes excrete their toxic agents after enzymatic neutralization is specific to normal liver tissue^13^ and has been reported to be remnant in some HB^9,10^.

**Fig. 3.**
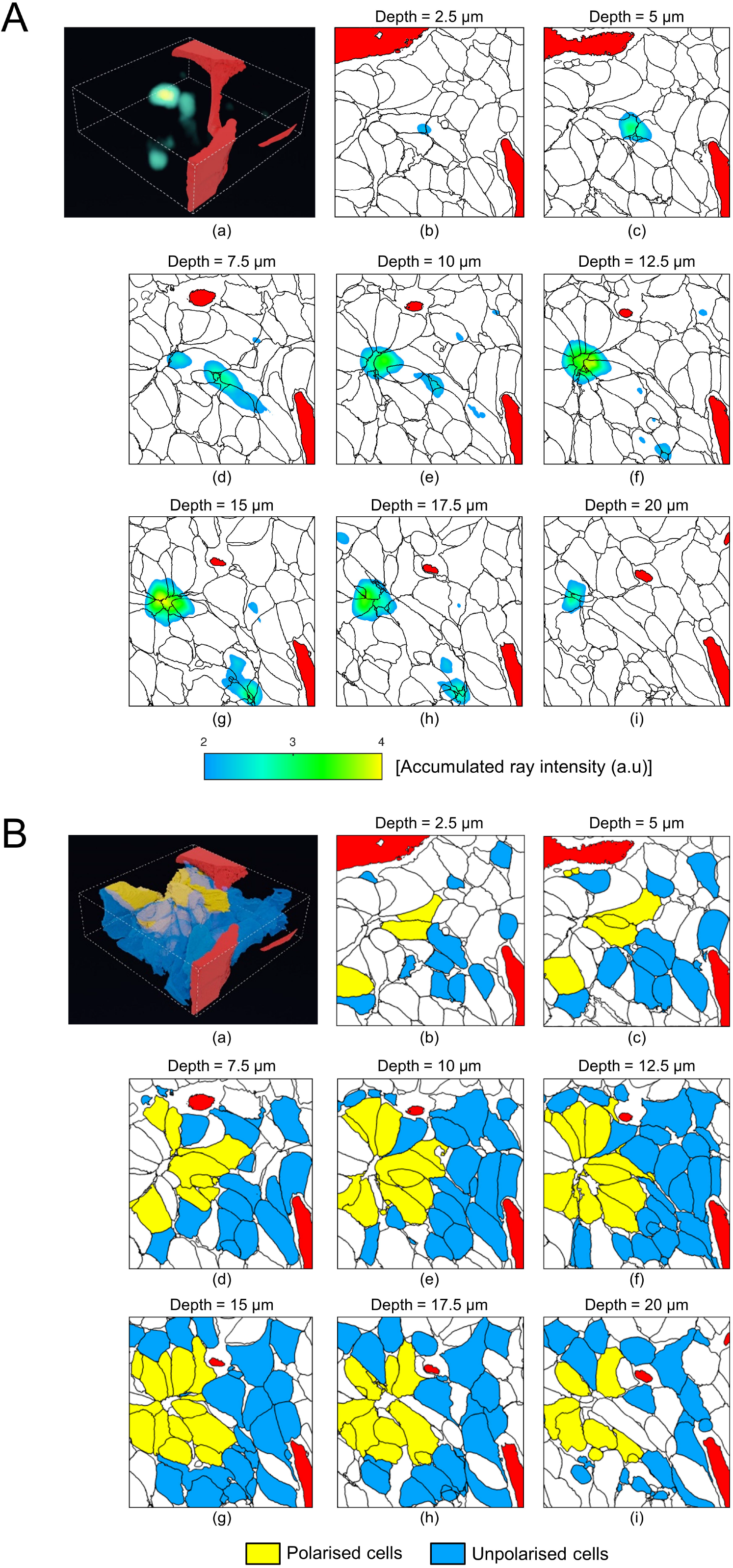
Tumour cell cluster with polarized shape orientation. (A-B) Only tumour cells with a complete nucleus are considered here. Blood capillary portions are in red. (A) Accumulation map of virtual rays [Accumulated ray intensity [a.u]; voxel-by-voxel basis; see yellowish voxels] emitted by each cell along its main axis. (b-i) 2D cross-sectional maps for increasing depth along the Z-axis. (B) Binary classification of polarised/unpolarised cells: 17 cells (in yellow) emitted virtual rays reaching the main accumulation region observable in panel A-(g) (accumulated ray intensity >3.5). a.u., arbitrary unit.

Finally, we focused our attention on the 21 fully segmented tumour cells contained in sample 2. Mitochondria were segmented by performing two successive cycles of automatic segmentation by deep-learning propagation combined with manual segmentation clean-up. Following the segmentation procedure, the size of these cells and their cytoplasm, nucleus and mitochondrial network was measured and component/cell ratios were calculated (Extended data Fig. 8A-B). In line with the known relationship between the size or number of organelles and the size of the cell^15,16^, we found that the larger the tumour cell, the larger its cytoplasm, global mitochondrial network and nucleus (Figure 4A). By measuring the distance between each of these cellular components and the blood capillary, we also found that the tumour cells located near the blood capillary were significantly larger than those located away from it (Fig. 4B-C, Supplementary Video 6). This inverse correlation with the distance to the capillary was also observed when considering the cytoplasm and the mitochondrial network (the latter having the highest correlative value) but not the nuclei (Fig. 4C, Extended data Fig. 9A-C, Supplementary Video 6). Altogether, these data suggested that the size of HB cells and their mitochondrial content is linked to the distance with blood capillary. These observations are in agreement with the decrease in mitochondrial mass in cells exposed to low oxygen supply^17,18^.

**Fig. 4.**
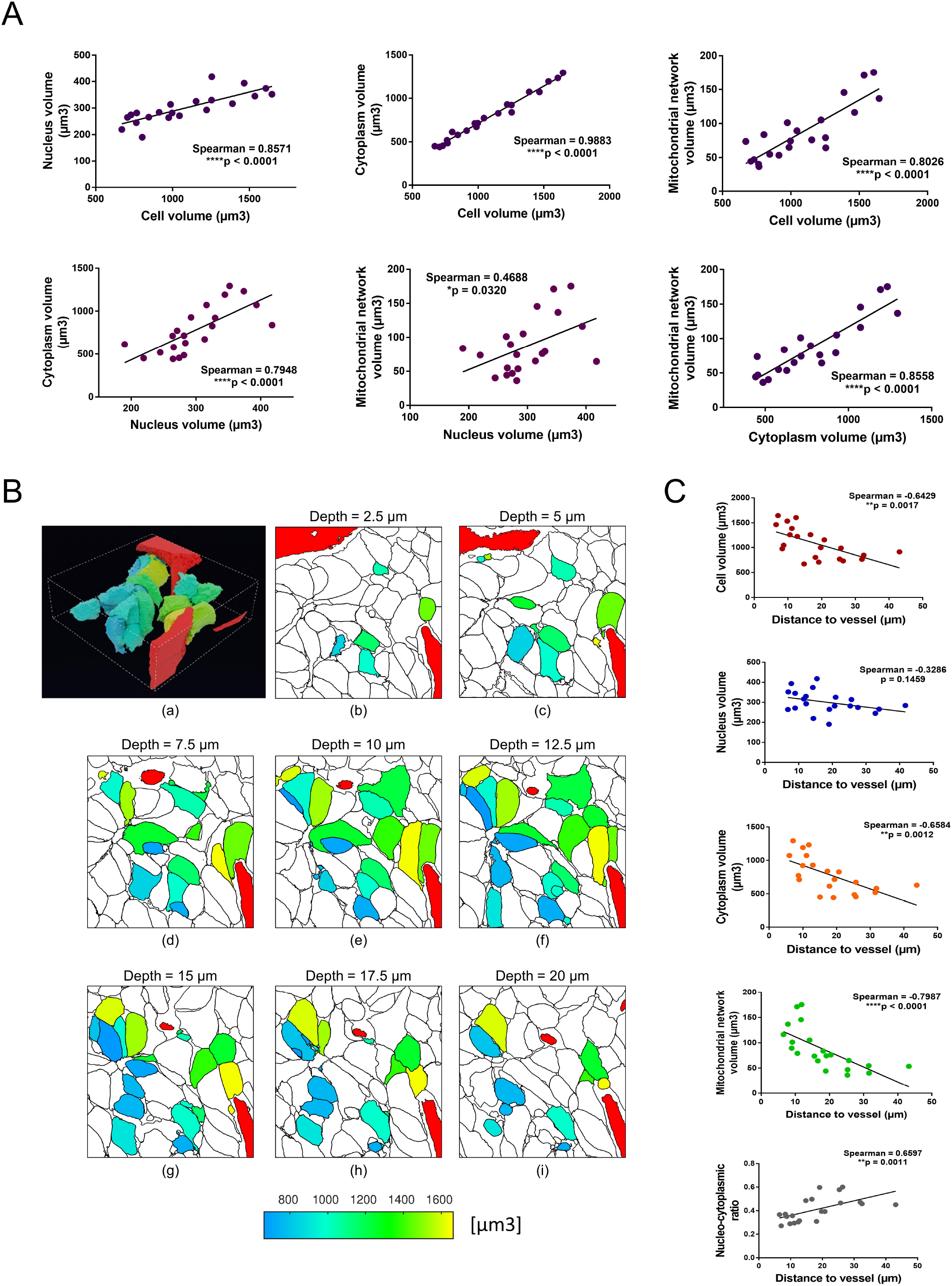
3D organization of typical cellular and subcellular structures. Only 21 entirely contained cells are considered here. (A) Volumetric correlations between cytoplasm, nucleus and mitochondrial network. (B) Cell sizes related to distance from blood capillary: (a) Reconstructed 3D image of cells and blood capillary. Cell colour is related to its volumetric size. (b-i) 2D cross-sectional maps for increasing depth along the Z-axis. (a-i) Blood capillary portions are in red. (C) Correlation between distance to blood capillary and cellular/subcellular structures or nucleocytoplasmic ratio. (A and C) Spearman correlations. r and p values are as indicated in corresponding graph.

## DISCUSSION

A still unanswered question is whether cells are distributed randomly in cancer tissue or if and how their organization is governed by physical and molecular factors reminiscent of the structure of normal tissue. In this pilot study, we used an onconanotomy approach, i.e. a workflow to analyse in depth the architecture of HB samples using a combination of sample preparation, SBF-SEM imaging, computational approaches and infographic tools. To our knowledge, this is the first report investigating the complete ultrastructure of human tumoral tissue by high-resolution 3D EM. Our data clearly demonstrate the feasibility of our methodology to study the 3D organization of tumoral tissue and the potential breakthrough it represents in understanding cancer biology. The information obtained on the organization of liver cancer cells throws much light on the physiology of tumours. In particular, it suggests that blood capillaries could determine the ultrastructure of tumoral tissue by controlling the alignment and size of tumour cells and their subcellular components, However, further studies are required to strengthen this hypothesis. In addition, we found that a bile canaliculus-like structure guides the spatial arrangement and the polarity of tumour cells in HB tissue, as it does in normal liver tissue to subsume the physiological functions of normal hepatocytes^13^.

This study is a preliminary attempt to unravel the internal architecture of tumoral tissue, so it has some limitations. First, it is difficult to automatically segment specific structures such as tumour-infiltrating macrophages and endoplasmic reticulum. The procedure requires visual control and manual correction, which are time-consuming. Second, our sample set was small. Future studies with a larger number and variety of tumoral tissues would allow us to generate more digitalized samples that in turn would improve the automatic segmentation process. A larger dataset would also allow us to better evaluate whether blood capillaries and bile canaliculus-like structures, which are common features in HB and more generally in epithelial liver tumours, influence the polarity and spatial organization of tumour cells and their organelles. These techniques coupled with current omic approaches would provide much insight into the relationship between the structural organization of tumours, and their cellular functions and metabolic activities.

Future possible directions of onconanotomy-based exploratory studies are the following: a) analysing virtually all types of solid tumours to generate a virtual 3D biorepository whose data would be particularly valuable in the case of rare tumours or precious samples such as biopsies; b) comparing matched tumour samples such as changes occurring before and after treatment or primary *versus* metastatic features, and c) identifying structural subtypes and verifying their overlap with different histological groups such as embryonal *versus* foetal *versus* small undifferentiated cell subtypes described in HB^6,14^. Since it is adapted to lab tumour models like cell xenografts, tumour-like spheroids and organoids, our combined approach could help investigators to clarify the role of genes and molecular pathways involved in the architecture of tumoral tissue and to investigate the response of tumour cells to drugs (e.g. checkpoint inhibitors, monoclonal antibodies) and treatments (e.g. radiotherapy).

In conclusion, our study aims to foster the development of what we call onconanotomy, a novel research field in oncology, that will complement other innovative imaging technologies such as two-photon excitation microscopy (also known as 2PEF) and correlative light-electron microscopy (also known as CLEM). This field could be backed up by the development of open-source databanks gathering all 3D imaging data at a worldwide level in order to boost research and promote cooperation in cancer, as was previously the case with Gene databanks and Cell and Tissue atlases. By accessing 3D image databanks, investigators could upload valuable information and perform comparative investigations to unravel the ultrastructure of solid tumours. The potential focuses include cell-cell and cell-extracellular matrix interactions, intratumoral heterogeneity, pro-metastatic mechanisms, angiogenesis, drug response, therapeutic efficacy and regulatory factor gradation. We believe that, in few decades from now and thanks to digital approaches and artificial intelligence, the emerging field of onconanotomy could profoundly change the face of medical oncology with the support of other technologies such as single-cell RNA-seq, microfluidics and organoids. It might especially provide ground-breaking insights into the predisposition of cells to disseminate and into the permeability of tissues to drugs. Overall, understanding the relationship between the organization of tumour cells and clinical outcome could help in identifying the key structural parameters of cancers. In turn, this could lead to major advances in our understanding of cancers and the most appropriate therapies to treat both adult and paediatric cancer patients.

## Acknowledgements

This work was supported by the charity Eva pour la Vie, La Fondation ARC pour la Recherche sur le Cancer (contract N° PJA 20191209631), La Région Nouvelle-Aquitaine, La Fondation Groupama pour la Santé and Groupama Centre-Atlantique. Microscopy Imaging was performed at the Bordeaux Imaging Centre, which is a member of the FranceBioImaging national infrastructure (ANR-10-INBS-04). Computational analysis was carried out using the PlaFRIM experimental testbed, supported by Inria, CNRS (LABRI and IMB), Université de Bordeaux, Bordeaux INP and Conseil Régional d’Aquitaine (see https://www.plafrim.fr/). Computer time for this study was provided by the computing facilities at MCIA (Mésocentre de Calcul Intensif Aquitain) of the Université de Bordeaux and of the Université de Pau et des Pays de l’Adour.

## Author contributions

Fresh paediatric tumour fragments were provided by S.B. and C.C. HB-PBX were developed by XenTech company and provided by K.F. and S.C. Image acquisition and analyses were performed by M.B., F.Z.K. and E.G. The manual segmentations and clean-up of tissue and cellular components were performed by F.Z.K., M.B. and C.F.G. B.D.d.S. performed the mathematical and computational analyses. A.L. generated 3D images and videos. C.F.G., S.C. and E.G. supervised the work. C.F.G. obtained the financial grants. All authors actively participated in the writing of the manuscript.

**Supplementary Information** is available for this paper.

## Abbreviations

2D: bi-dimensional
3D: tri-dimensional
CAM: chick embryo chorioallantoic membrane
EM: electron microscopy
HB: hepatoblastoma
PCA: principal component analysis
PDX: patient-derived xenograft
ROI: region of interest
SBF: serial block-face
SD: standard deviation
SEM: scanning electron microscopy
TEM: transmission electron microscopy

## METHODS

### PDX samples

PDX were generated in compliance with the informed consent form signed by the patients and developed as previously described ^7^. The main clinical and genetic characteristics of HB PDX samples are shown in Extended Table 1.

### Serial block-face scanning electron microscopy sample preparation

Tissue was prepared for SBF-SEM as previously described ^8^. Once the resin block was hardened, it was cut roughly with a razor blade to create a pyramid shape and mounted on aluminium specimen pins using a silver-filled conductive resin (Epotek-Delta Microscopies, Mauressac - France). After 24hr of polymerisation at 60°C, the samples were trimmed with a diamond knife (Diatome – Nidau - Switzerland). Silver-filled conductive resin was used to connect the edges of the tissue electrically to the aluminium pin. The entire sample was then sputter-coated with a 5-10 nm layer of gold to enhance conductivity.

### Transmission Electron Microscopy

The samples were first analysed by TEM to control morphology and define the region of interest. Ultra-thin sections were cut and deposited on copper grids and observed with a Hitachi H7650 transmission electron microscope (Japan).

### Serial block-face scanning electron microscopy imaging

The sample on the pin was placed into the carrier that fits into the3View ultramicrotome (3viewXP2-Gatan Inc., Pleasanton, CA, USA) on a ZEISS Gemini field emission gun SEM300 (Zeiss - Marly-le-Roi - France). The block face was imaged with an accelerating voltage at 1.2kV using the Back scattered electron with a specific BSE Detector (On point - Gatan Inc., Pleasanton, CA, USA).

### Pre-processing and structure segmentation

Image analysis was performed using three complementary software packages. Digital Micrograph (Gatan Inc., Pleasanton, CA, USA) was used to align the images with the Image Alignment plugin using the combined default filter (which combines soft rectangle and band pass filters). The aligned images were saved in Gatan format “dm4” as a single stack. Fiji software was used to convert the 32-Bit images into 8-Bit images to be less resource-consuming and facilitate segmentation. Then the brightness and contrast were automatically adjusted. Finally, VAST-Lite software was used for the manual segmentation of the elements of interest (cells, nuclei and mitochondria) ^12^. All manual segmentation was done using an interactive drawing tablet and pen (Cintiq pro 5, Wacom). The manual segmentation of both cells and nuclei was done only on 10% of the stack of images (1 image out of 10). These manual segmentations were propagated to neighbouring slices using a so-called « Optical Flow » algorithm (developed under the commercial software Matlab ©1994-2021 The MathWorks, Inc.) applied on the EM images, similar to the approach described by Huang T.C. and al ^19^. Mitochondria, which are much contrasted structures, were segmented manually on one single cell to feed a deep-learning algorithm (using a 2D U-net architecture ^20^ and an implementation based on Tensorflow 1.4 and Keras 2.2.4), which could be subsequently used for semi-automatic segmentation. Note that all semi-automatic segmentation results underwent human control and a manual clean-up was done when needed.

### Analysis of orientation of structures

Each cell, nucleus and capillary portion was characterized using its main axis. To this end, a Principal Component Analysis (PCA) was applied on the voxel coordinates within the segmented mask of a structure of interest. Only tumour cells with a complete nucleus were considered. The following two analyses were conducted.

#### Analysis of alignment in 2D plane of cells and nuclei

The main axes of cells were used to calculate the best 2D “alignment plane” (in the least-squares sense: we minimized the sum of squared differences between the observed main axes of cells, and the fitted value provided by a 2D plane equation). Angles between the main axis of each cell and the alignment plane were calculated. The same analysis was conducted subsequently for nuclei.

#### Determination of cell clusters with polarized shape orientation

3D virtual rays were emitted by each cell along its main axis in both directions (the ray radius was a user-defined input parameter for the algorithm and was set to 15 μm, ray intensity proportional to the Euclidean distance to the main axis, with a value of 1 on the main axis). To this end, Bresenham’s line algorithm ^21^ was employed and adapted to our need. The accumulated beam intensity was calculated on a voxel-by-voxel basis. Cells emitting virtual rays reaching a specific accumulation region were subsequently selected. Thus, cells converging toward a common region could be listed.

### Infographics

Blender version 2.90.1 (https://www.blender.org) was used to create the illustrations (images and Supplementary videos). The Cycles render engine was used for all the final renderings. To render polygonal meshes, the data were imported from VTK files using the BVTK addon (https://github.com/tkeskita/BVtkNodes). Depending on the data complexity, the meshes were decimated to lower the polygon count, but still preserving the overall shape and most of the details. For volumetric rendering, the original stack data was converted from a multi-page Tiff image file to a downscaled 2D tile map, which was then loaded and translated to a 3D volume into Blender using a custom shader setup. ImageJ (https://imagej.nih.gov/ij/), XNview (https://www.xnview.com) and the “montage” command line tool of ImageMagick (https://imagemagick.org/script/montage.php) were used for image conversion, downscaling and stitching. Downscaling is mandatory, since the original amount of data is not always practical to work with and is not necessary for the purpose of visualization.

### Statistical analyses

Statistical analyses were performed using GraphPad Prism 7.05 software. Spearman’s nonparametric correlation was used to compare the size of different cell components or the size of a specific component to the distance to a blood capillary. Results were considered significant when p < 0.05.

## Supplementary Information Guide

**Supplementary Video 1: Spatial organization of blood capillary portions and circulating cells in HB PDX Sample 1.** See also legend of Extended data Fig.2.

**Supplementary Video 2: 3D organisation of tumour cell and its main subcellular components in HB PDX Sample 2.** See legend of Extended data Fig.3.

**Supplementary Video 3: Tumour and non-tumour components of HB PDX Sample 2.** See legend of Fig. 2A-C.

**Supplementary Video 4: Alignment in plane of tumour cells or tumour cell nuclei *versus* blood capillary in HB PDX Sample 2.** See legend of Fig. 2D.

**Supplementary Video 5: A set of tumour cells polarize in direction of a bile canaliculus-like structure.** See legend of Fig. 3.

**Supplementary Video 6: Spatial organization of 21 cells and their organelles in HB PDX Sample 2.** See legends of Fig. 4B and of Extended data Fig. 9.

