## Supplementary figures and images for "Deciphering Tumour Tissue Organization by 3D Electron Microscopy and machine learning"

### Extended Data Fig.1

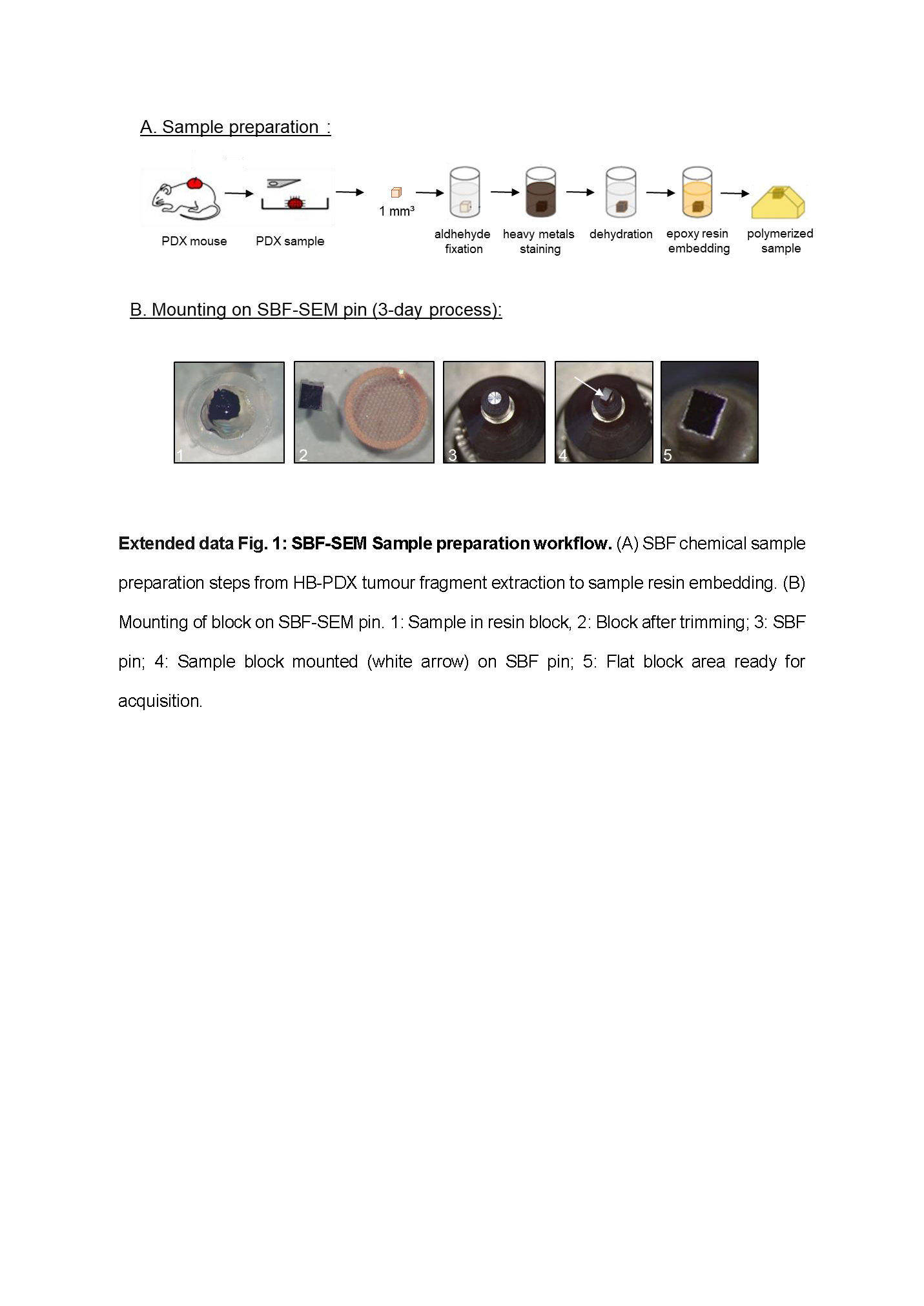

### Extended Data Fig.2

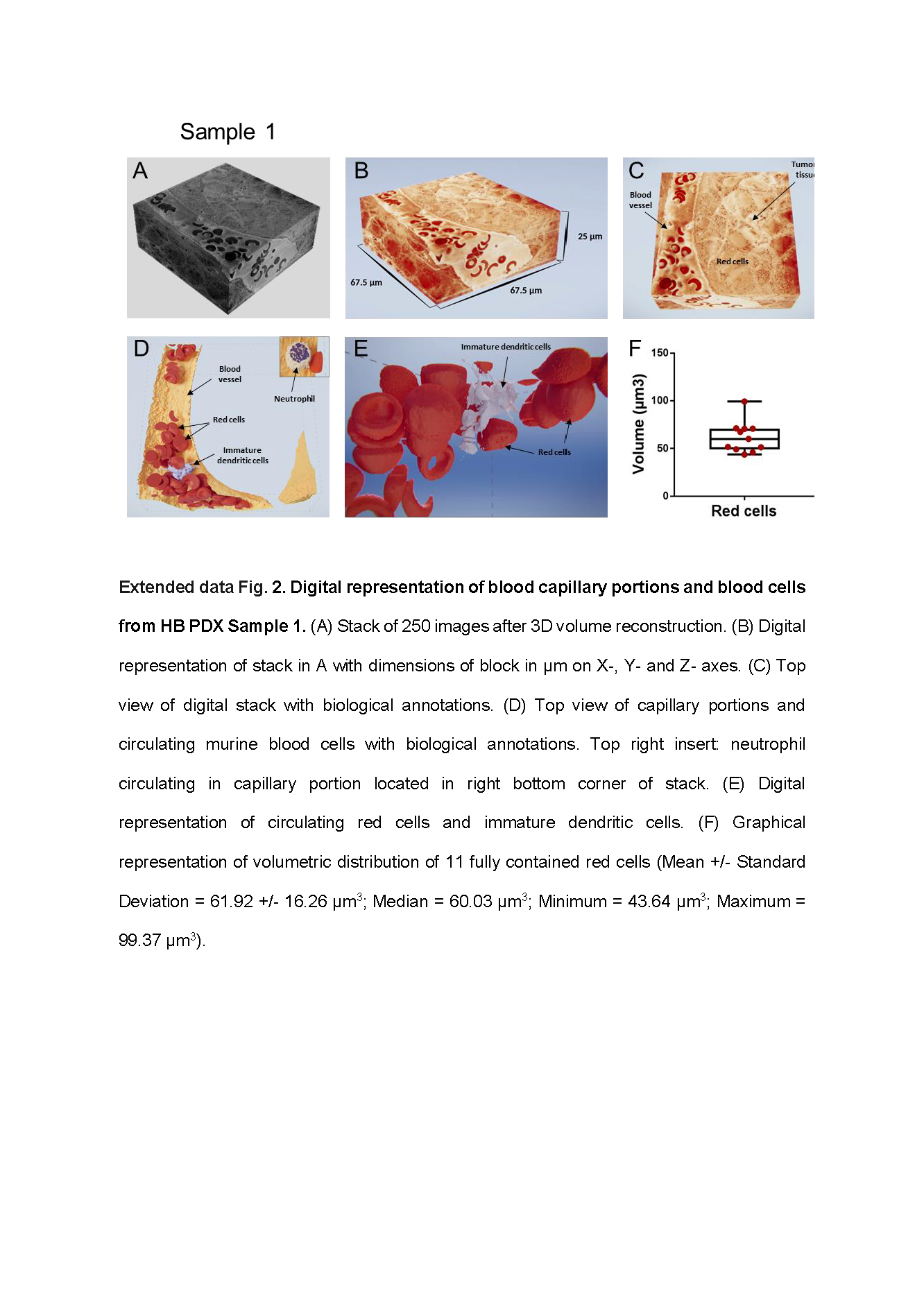

### Extended Data Fig.3

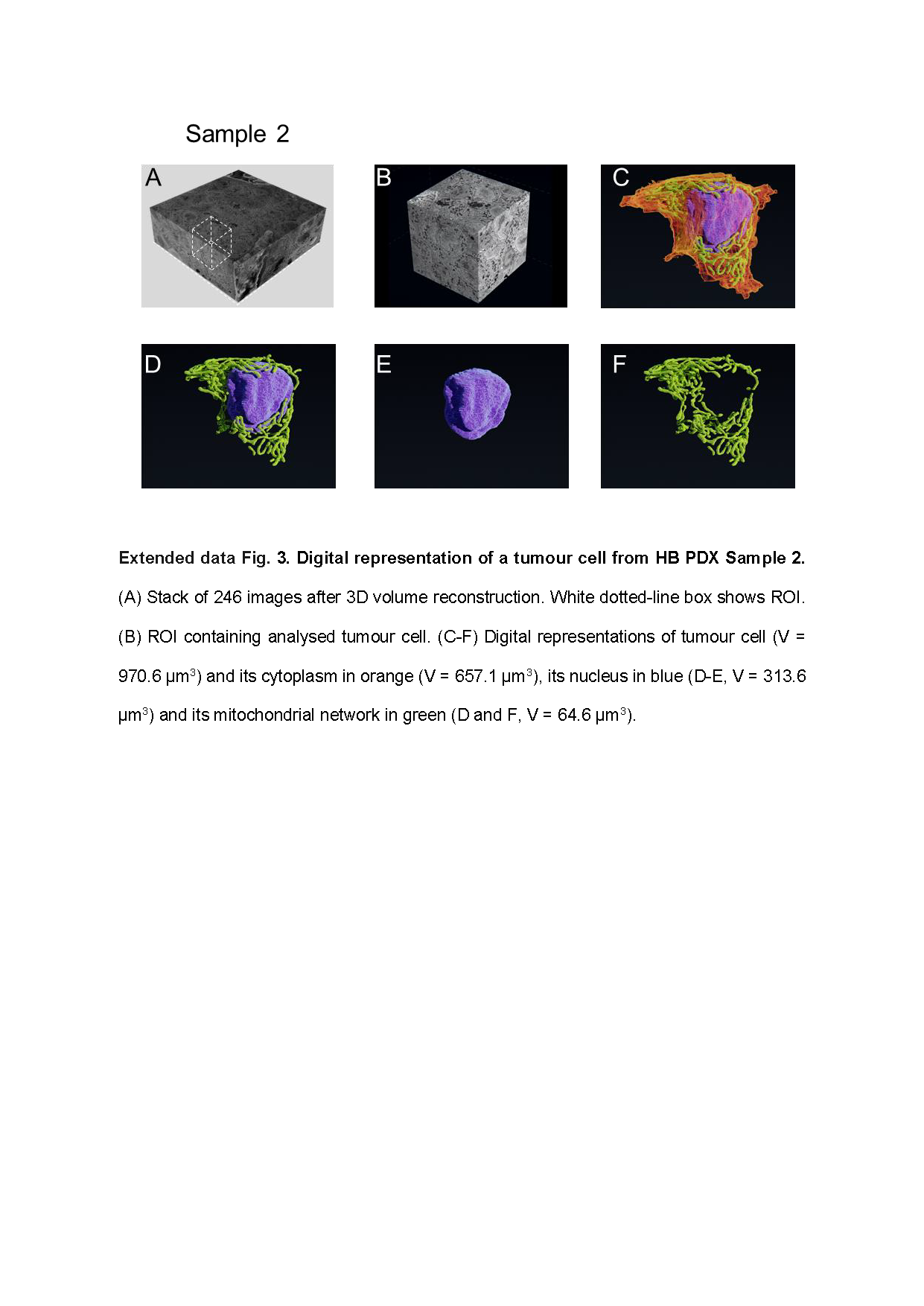

### Extended Data Fig.4

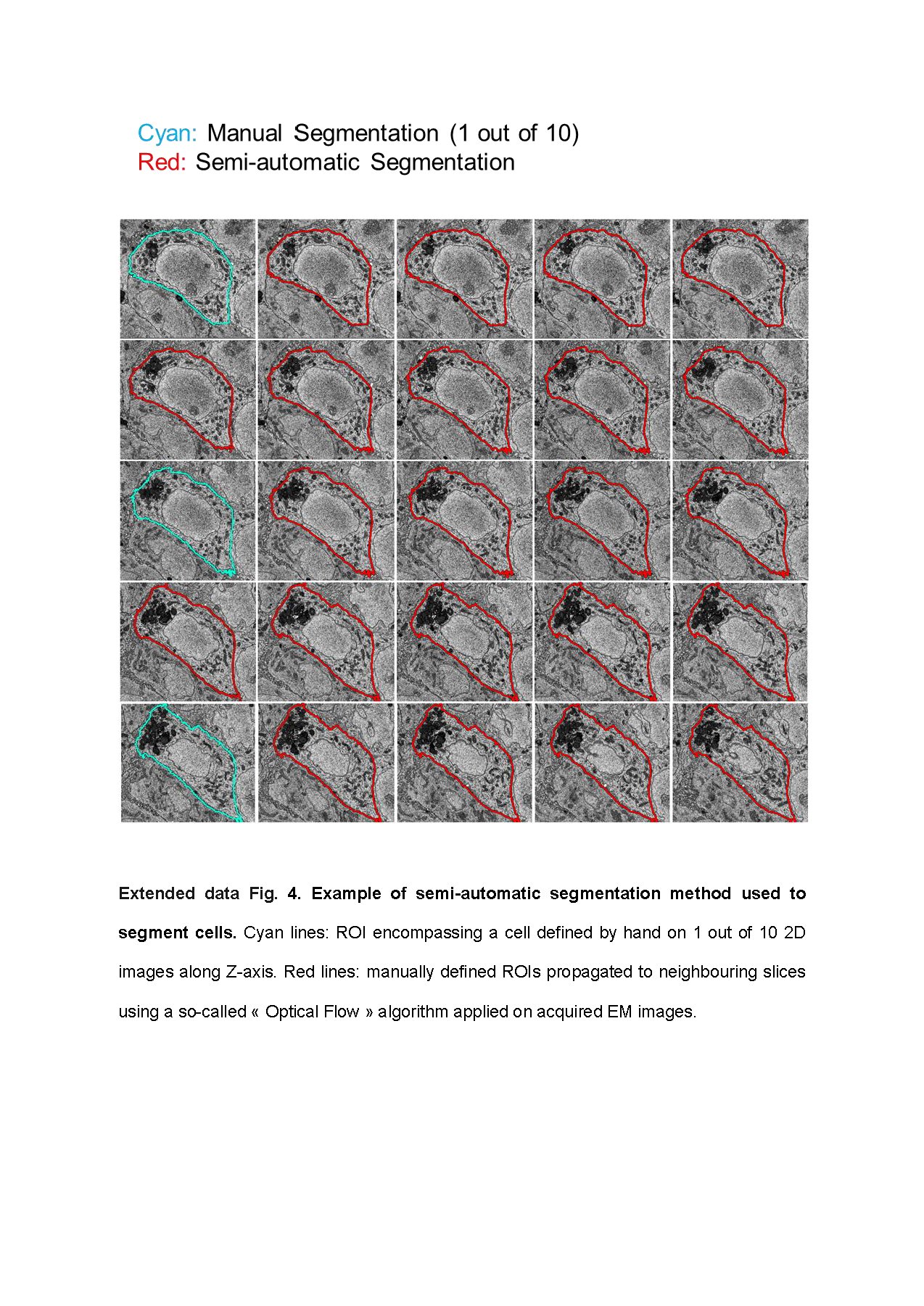

### Extended Data Fig.5

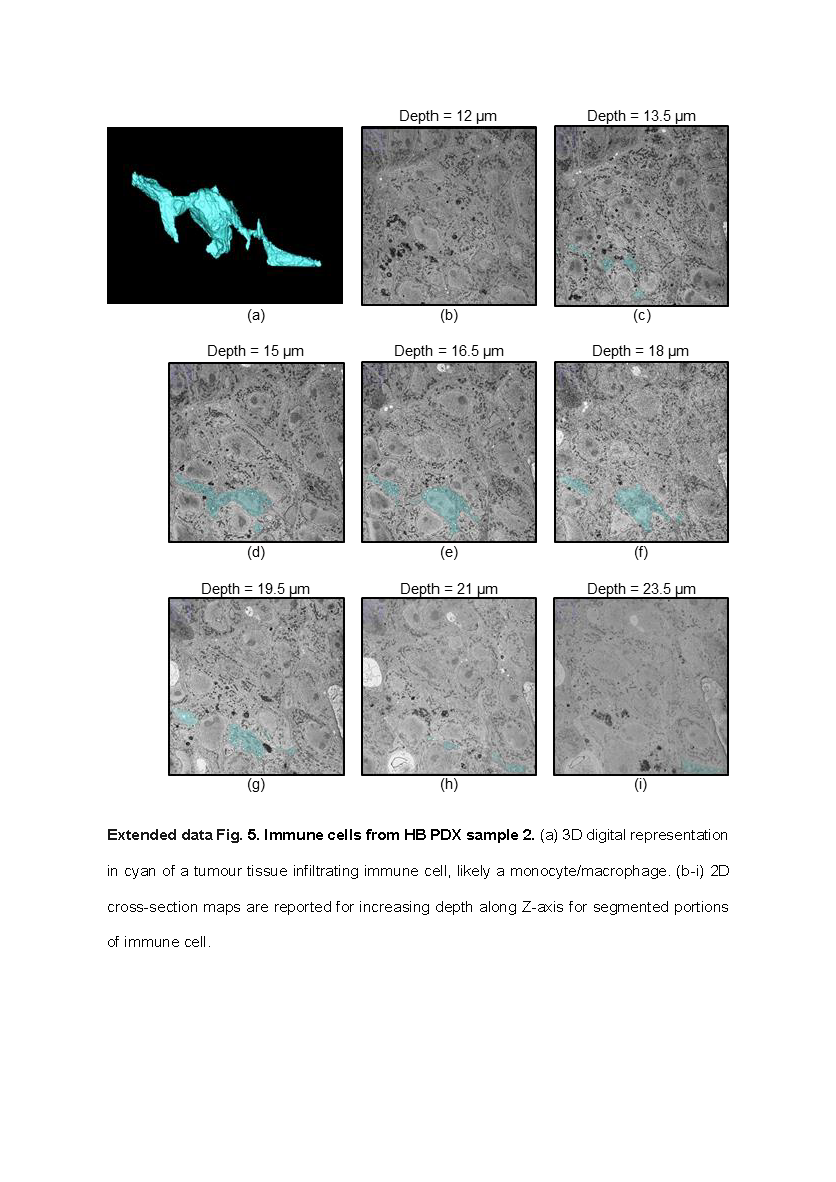

### Extended Data Fig.6

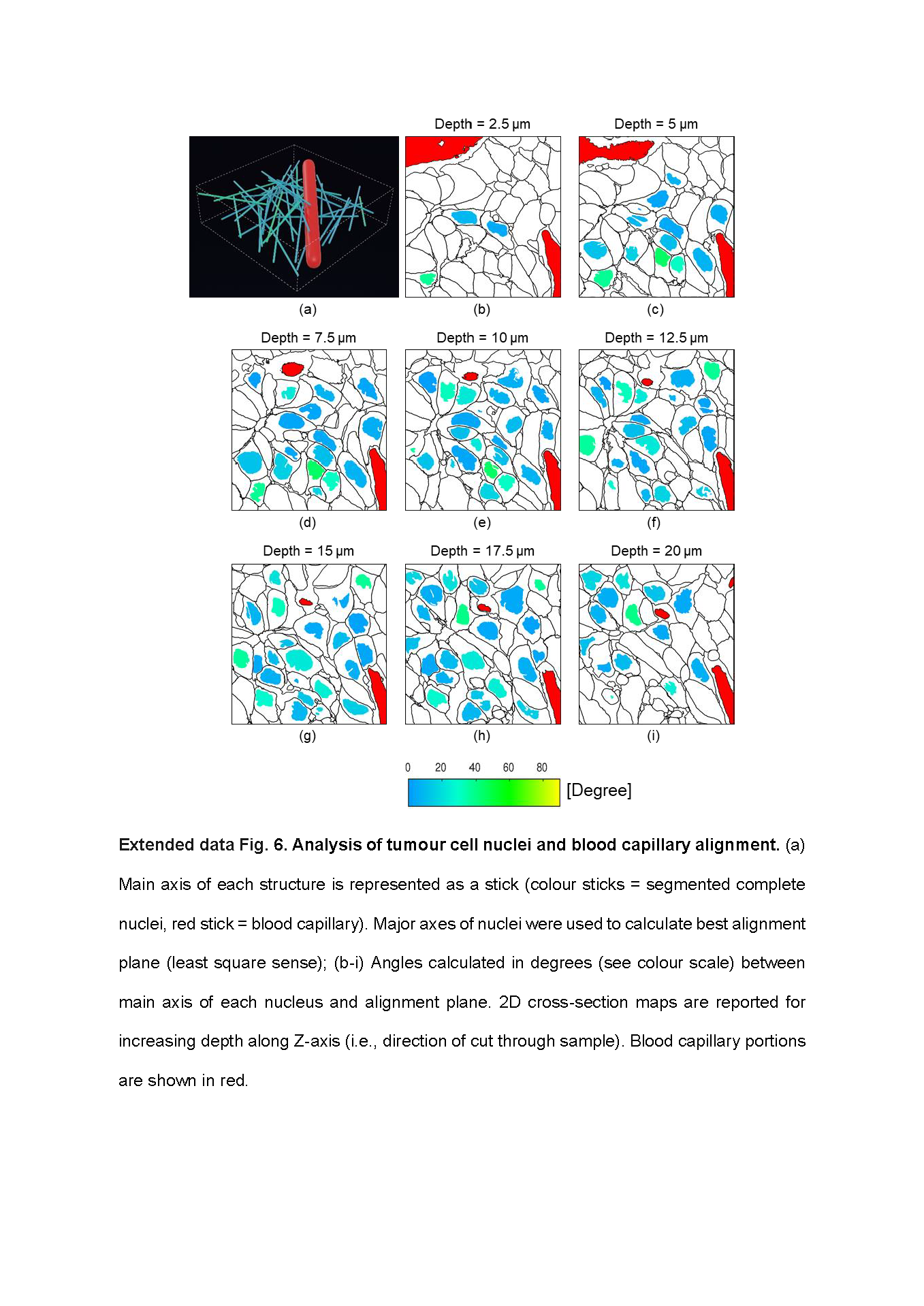

### Extended Data Fig.7

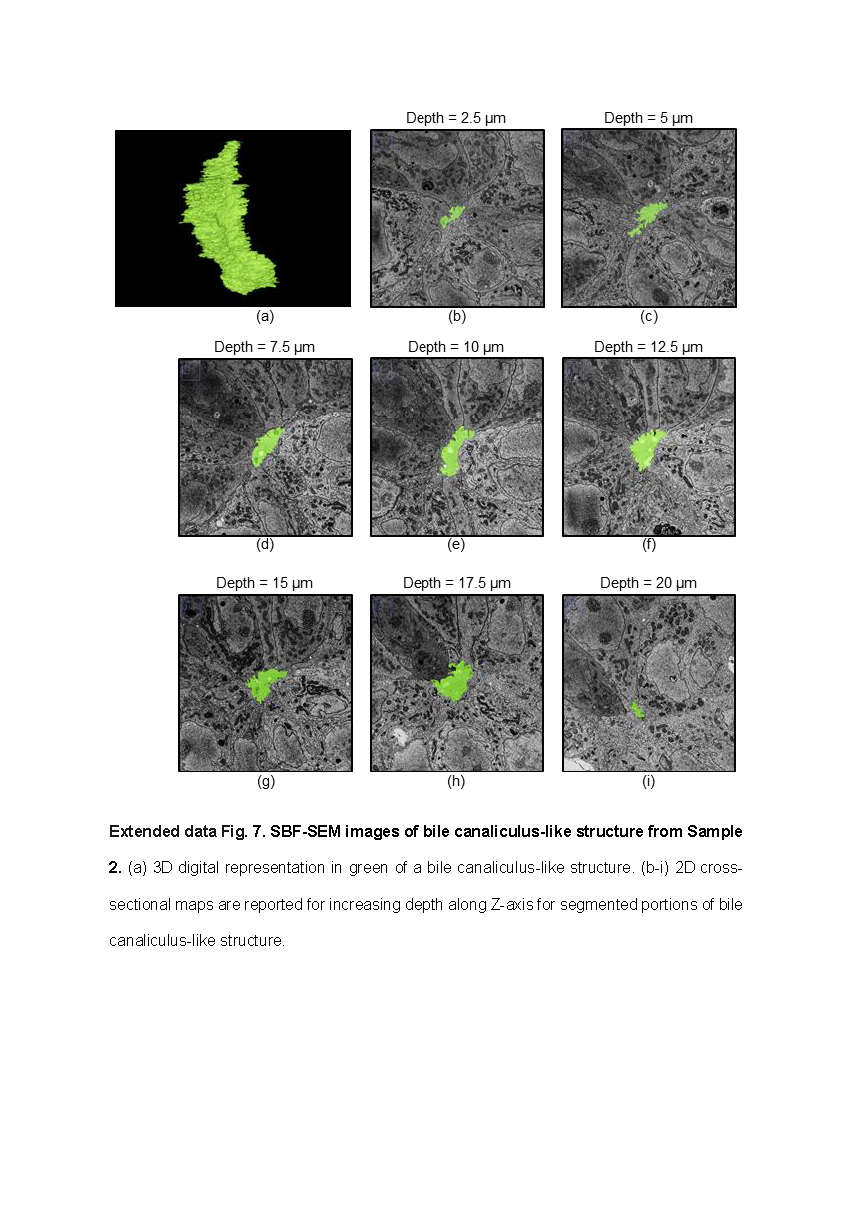

### Extended Data Fig.8

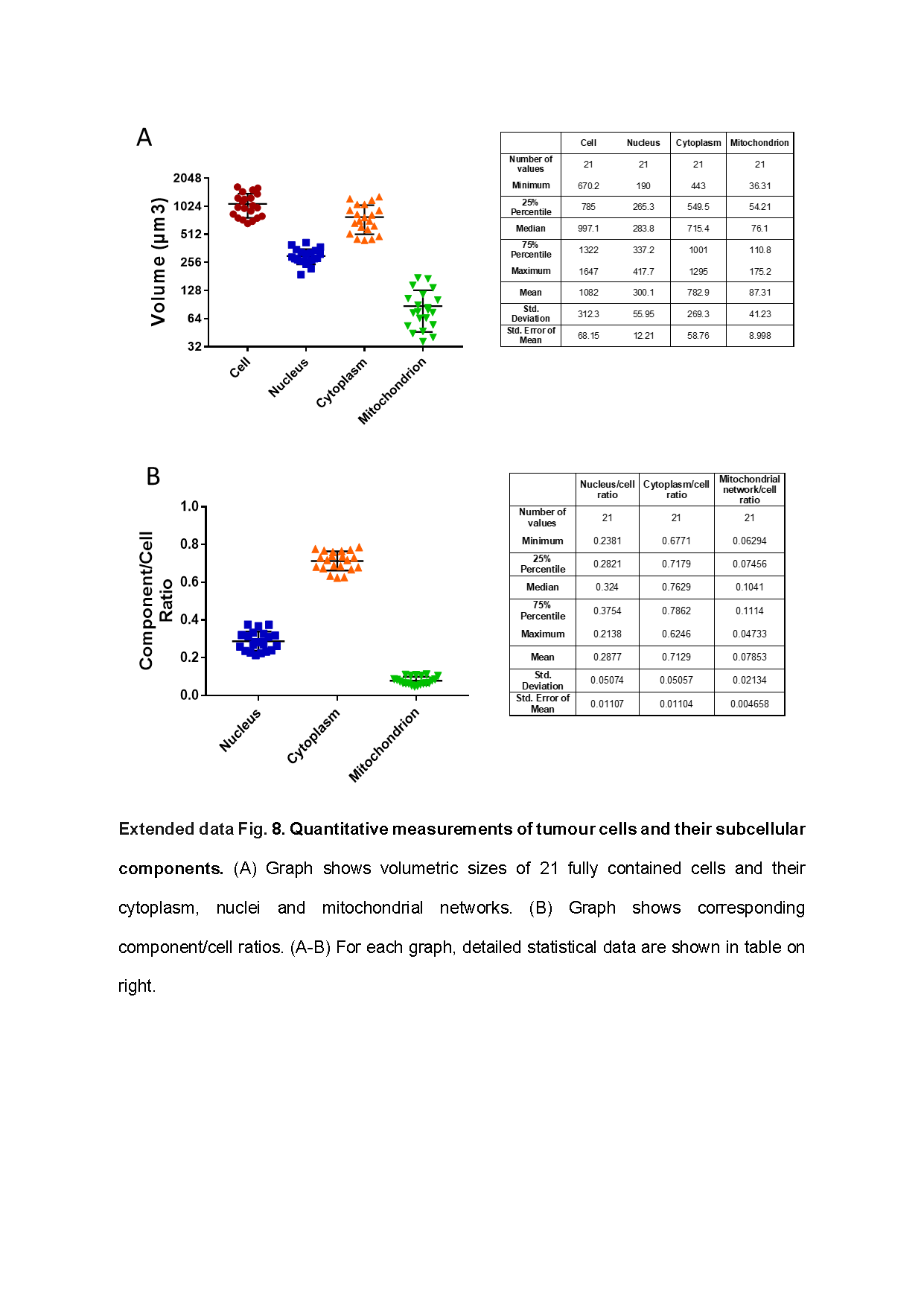

### Extended Data Fig.9

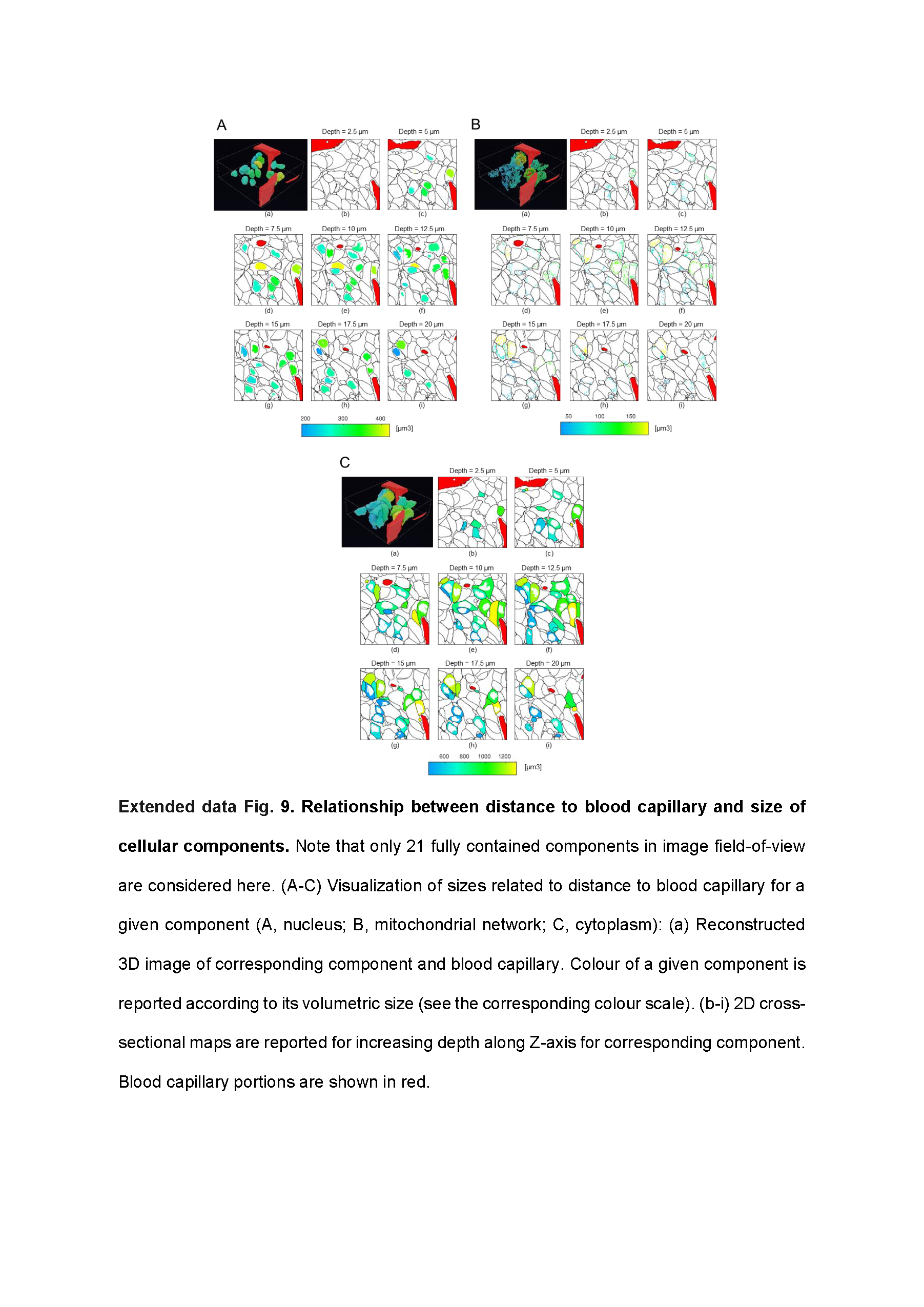

### Extended Data Table 1

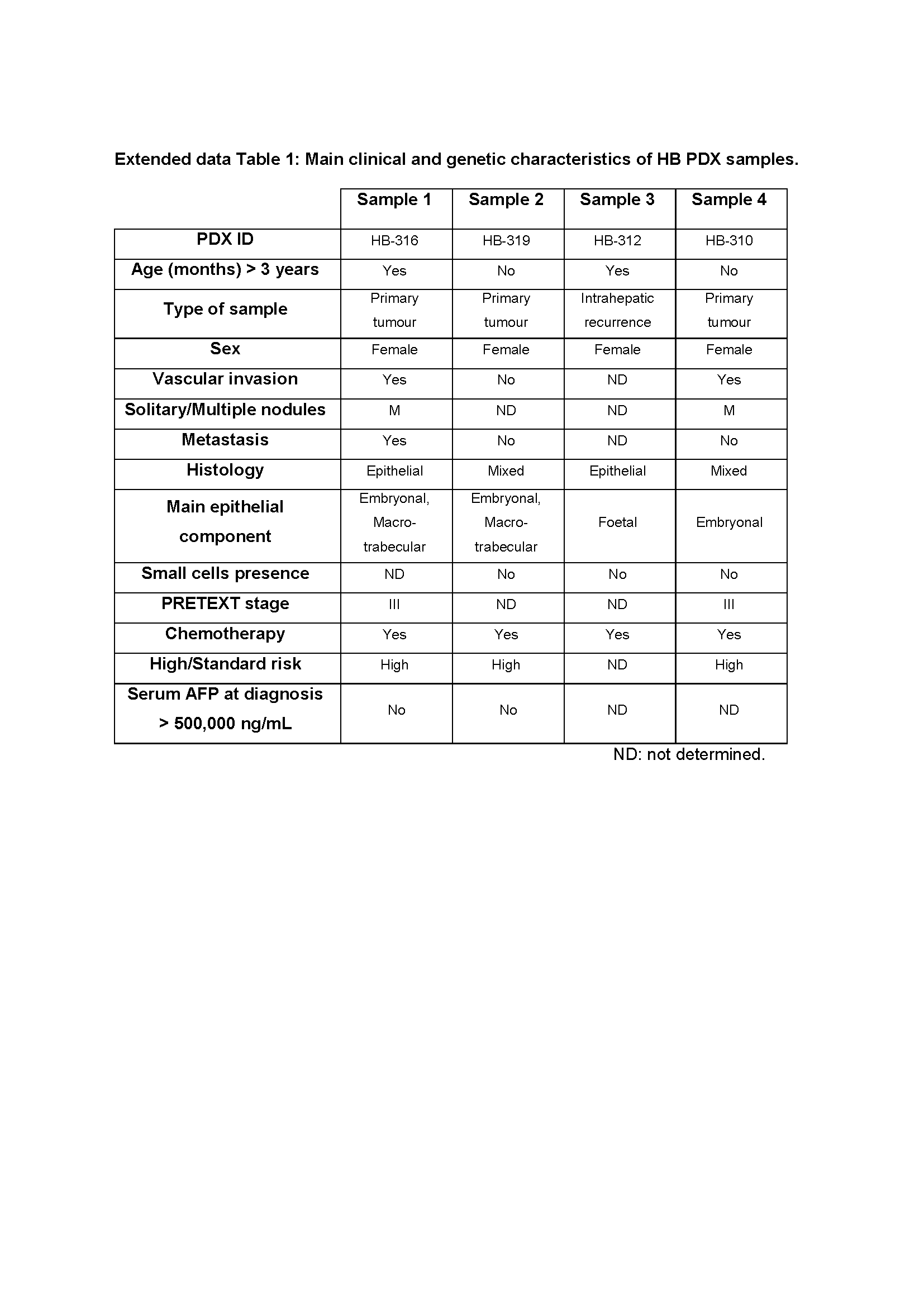
